# A series of constitutive expression vectors to accurately measure the rate of DNA transposition and correct for auto-inhibition

**DOI:** 10.1101/423012

**Authors:** Michael Tellier, Ronald Chalmers

## Abstract

**Background:** Transposable elements (TEs) form a diverse group of DNA sequences encoding functions for their own mobility. This ability has been exploited as a powerful tool for molecular biology and genomics techniques. However, their use is sometimes limited because their activity is auto-regulated to allow them to cohabit within their hosts without causing excessive genomic damage. To overcome these limitations, it is important to develop efficient and simple screening assays for hyperactive transposases.

**Results:** To widen the range of transposase expression normally accessible with inducible promoters, we have constructed a set of vectors based on constitutive promoters of different strengths. We characterized and validated our expression vectors with Hsmar1, a member of the *mariner* transposon family. We observed the highest rate of transposition with the weakest promoters. We went on to investigate the effects of mutations in the Hsmar1 transposase dimer interface and of covalently linking two transposase monomers in a single-chain dimer. We also tested the severity of mutations in the lineage leading to the human *SETMAR* gene, in which one copy of the Hsmar1 transposase has contributed a domain.

**Conclusions:** We generated a set of vectors to provide a wide range of transposase expression which will be useful for screening libraries of transposase mutants. We also found that mutations in the Hsmar1 dimer interface provides resistance to overproduction inhibition in bacteria, which could be valuable for improving bacterial transposon mutagenesis techniques.

## Background

Transposable elements (TEs) are DNA sequences encoding their own ability to move in a genome from one place to another. They are found in virtually all organisms and are particularly present in eukaryotes where they can represent a high percentage of the genome (1-3). Originally described as selfish elements since they were considered parasites which use the host for propagation but do not provide any particular advantage, TEs have now been shown to be important drivers of genome evolution (4, 5). Indeed, TEs can provide novel transcription factor binding sites, promoters, exons or poly(A) sites and can also be co-opted as microRNAs or long intergenic RNAs (6-8). TEs are a diverse group of DNA sequences using a wide range of mechanisms to transpose within their hosts. One particular mechanism prevalent in eukaryotes, and used by the *mariner* family, is known as “cut-and-paste” transposition (9). Over the past several years, our group and others have described the mechanisms regulating the transposition rate of different *mariner* transposons, such as Himar1, Hsmar1 or Mos1 (10-15). In Hsmar1, a regulatory mechanism was first recognized because of the phenomenon of overproduction inhibition (OPI) (16). The mechanism of OPI was eventually explained by the realization that double occupancy of the transposon ends with transposase dimers blocks assembly of the transpososome (12). Thus, OPI curbs Hsmar1 transposition rate to avoid damaging the host genome by excessive transposition (12).

However, OPI represents a limitation in the development of hyperactive transposases, which would facilitate transposon mutagenesis. Several approaches such as modifying the binding kinetics of the transposase to the inverted terminal repeat (ITR) or the monomer-dimer equilibrium can be used to overcome OPI. Indeed, we and others previously showed that most mutations in the conserved motif, WVPHEL, in Himar1 and Hsmar1, located at the subunit interface, result in hyperactive transposases but at the cost of producing non-productive DNA double-strand breaks and therefore DNA damage (17, 18).

To facilitate the isolation of suitable transposase mutants, the papillation assay was developed as an efficient screening procedure (Supplementary Figure 1) (19, 20). This assay is based on a promoter-less *lacZ* gene flanked by transposon ends. This reporter is integrated in a silent region of the genome of *Escherichia coli*. The transposase gene is provided *in trans* on a plasmid to simplify mutagenesis and library handling. Transposition events into an expressed ORF give rise to lacZ gene fusion proteins. When this happens within a colony growing on an X-gal indicator plate, it converts the cell to a lac+ phenotype, which allows the outgrowth of blue microcolonies (papillae) on a background of white cells. The transposition rate is estimated by the number of papillae per colony and by the rate of their appearance.

A limitation of the papillation assay is that it generally employs a transposase gene whose expression is under the control of an inducible promoter which cannot be finely regulated. We have constructed a set of vectors maintained in single copy or as five copies per cell which carry various constitutive promoters in the absence or presence of a ribosome binding site (RBS). This set of vectors allows transposase expression across a wide range of expression levels facilitating the screening of hyperactive and/or OPI-resistant transposases. We used this set of vectors to compare an Hsmar1 transposase monomer to a single-chain dimer and to test for hyperactivity and OPI-resistance several Hsmar1 transposase mutants. We found that one Hsmar1 mutant in the dimer interface, R141L, is resistant to OPI in *E. coli*.

## Results and Discussion

### Characterization of the papillation assay using a strong inducible promoter

The papillation assay provides a visual assessment of the transposition rate, which can be determined from the rate of papillae appearance and their number per colony (19). The transposition rate is dependent on the concentration and activity of the transposase (12). We defined the transposition rate as the average number of papillae per colony after five days of incubation at 37°C. In the existing papillation assay, the transposase was provided by the protein expression vector pMAL-c2x under the control of a Ptac promoter (18). We first characterized the papillation assay using the Hsmar1 transposase cloned downstream of the inducible Ptac promoter and investigated the effect of different concentrations of IPTG and lactose and the presence or absence of the MBP tag on the transposition rate (Figure 1). In absence of transposase, the number of papillae per colony in all the conditions tested is either zero or one (Figure 1, Ø lane). In presence of the transposase or MBP-transposase (middle and right lanes, respectively), the number of papillae per colony varies with the concentration of IPTG and lactose.

**Figure 1.**
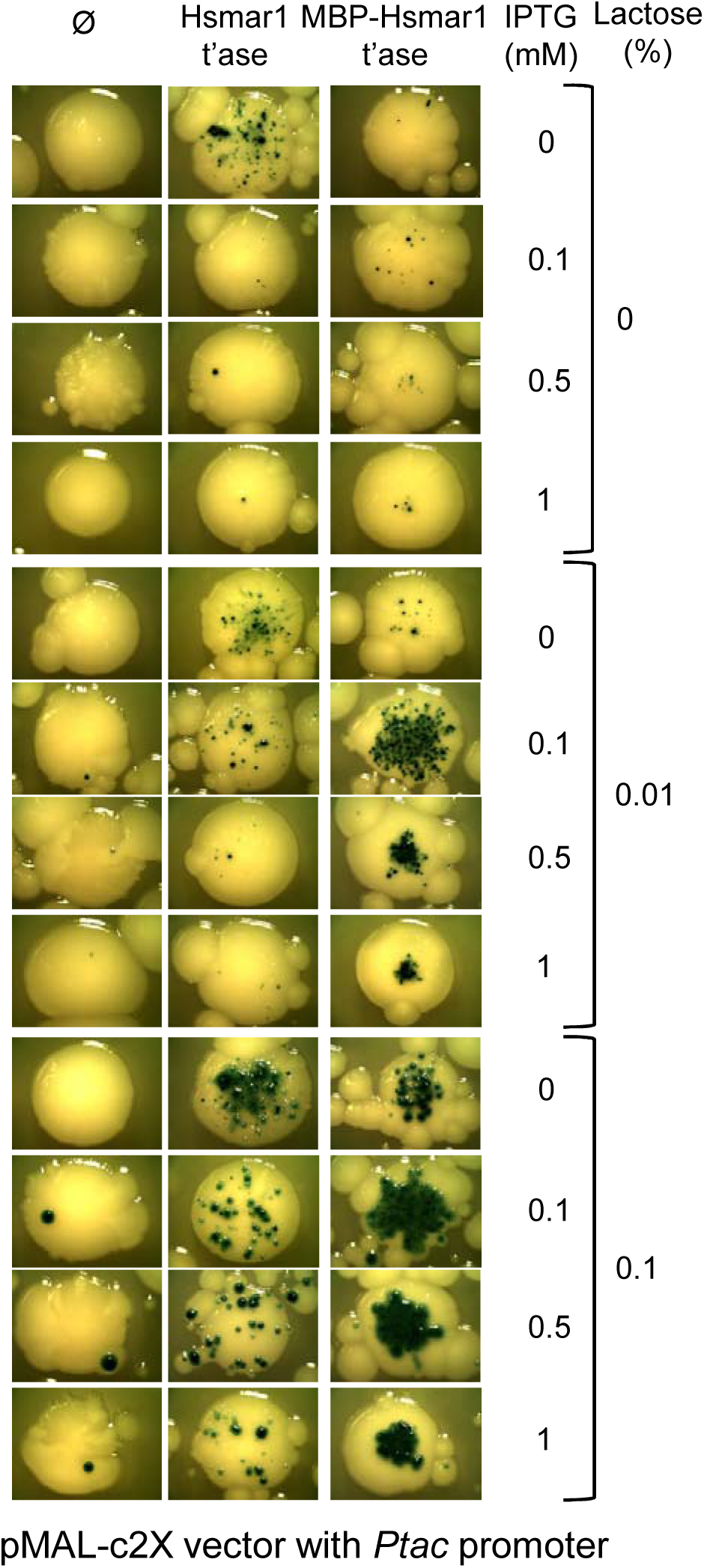
Characterization of the papillation assay using a strong inducible promoter. An expression vector encoding Hsmar1 (pRC1721) or MBP-Hsmar1 (pRC880) transposase (t’ase) was transformed into the papillation strain and plated on different lactose and IPTG concentrations. Representative colonies of the papillation plates are shown. On some pictures, smaller colonies surrounding the main colony are visible. These satellite colonies appear only after several days of incubation when the ampicillin present on the plate has been degraded. They can be ignored because they do not contain any transposase expression plasmid. Part of this figure was previously published in (21).

Independently of the presence or absence of the MBP tag and the IPTG concentration, the number of papillae increases with the concentration of lactose (Figure 1). Lactose improves the sensitivity of the assay by allowing papillae to continue to grow when non-lactose carbon sources are exhausted. At all lactose concentrations, the transposition rate is the highest at 0 and 0.1 mM IPTG for the transposase and the MBP-transposase, respectively (Figure 1). Any further increase in the IPTG concentration results in a decrease of the transposition rate, consistent with the effects of overproduction inhibition (OPI), which has been described for Hsmar1 *in vitro*, in *E. coli*, and in HeLa cells (12, 21). Interestingly, the presence of the MBP tag appears to lower the transpositional potential of the system, potentially through the stabilization of the Hsmar1 transposase. We therefore decided to use untagged Hsmar1 transposase for the remaining experiments.

### Papillation assay with a featureless DNA constitutive promoter

We wondered if the expression level of the un-tagged transposase at 0 mM IPTG (Figure 1) represents the peak activity of the system or is the system already in OPI? To answer this question, we took advantage of a 44 GACT repeats sequence that represents an idealized segment of unbent, featureless DNA. It is known as the “even end” (EE) as it was first used to study the role of DNA bending in Tn10 transposition (22). We reasoned that this would provide for a minimal level of transcription owing to its lack of TA and AT dinucleotides that feature in the -10 region of sigma70 promoters (Figure 2A, RBS^+^). Although the EE does not provide a -10 region, it provides a G+A rich sequence for ribosome binding. We therefore abolished or optimized this putative RBS (Figure 2A, RBS^-^ and RBS^++^, respectively). We find that transposition is the highest in absence of a RBS (Figure 2B and C).

**Figure 2.**
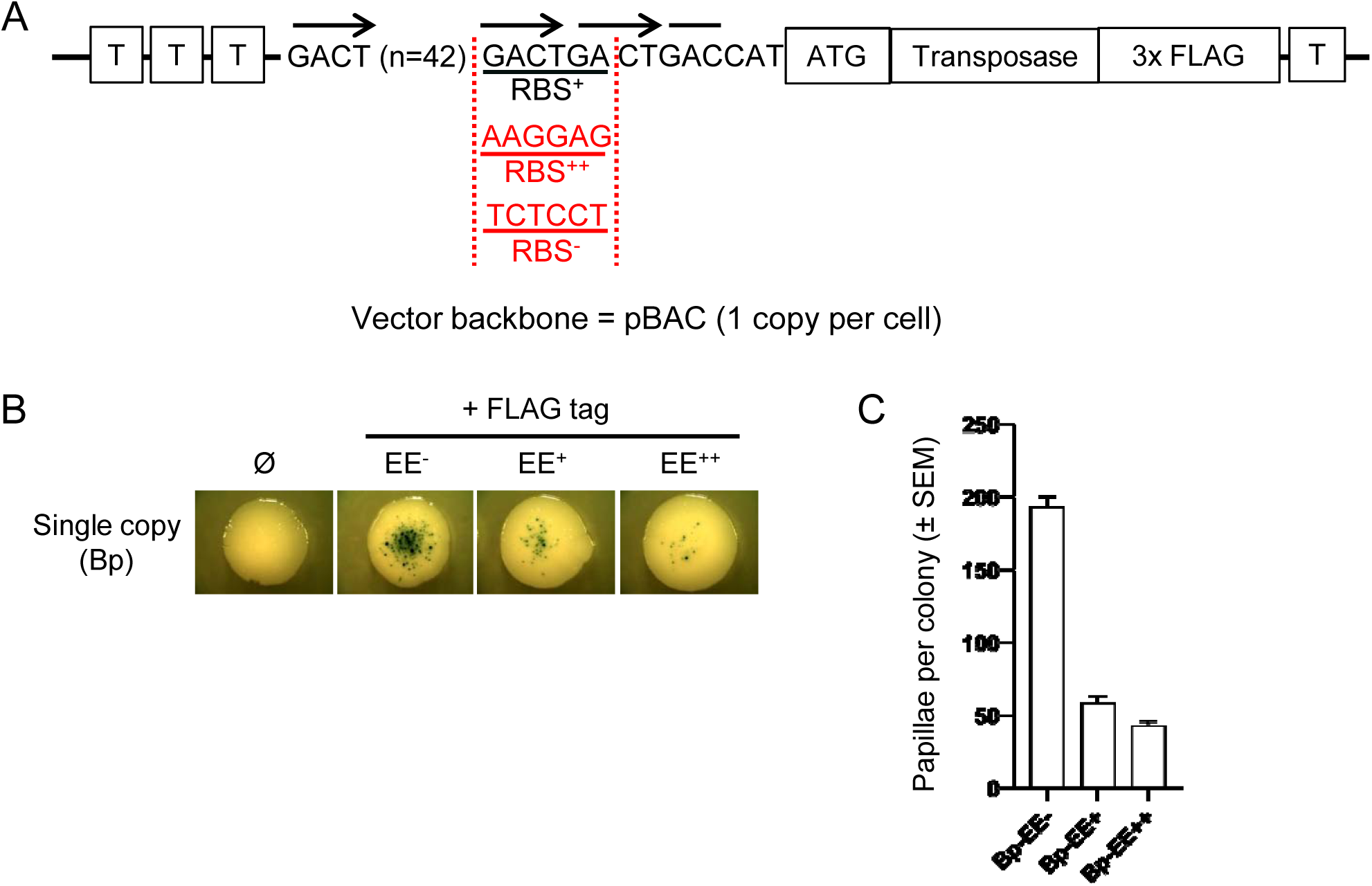
Papillation assay with a featureless DNA constitutive promoter. **A/** The *Hsmar1* gene is fused to 3x Flag tag on its C-terminus and cloned downstream of pEE containing a ribosome binding site (RBS) based on the GACT repeat (RBS+), on an optimal RBS sequence (RBS++), or on an inactive RBS sequence (RBS-). The construct is located between terminator sequences (T) upstream and downstream to avoid read-through transcription. The plasmid backbone is a one-copy vector, pBACe3.6. **B/** Representative colonies of each single-copy vector expressing a wild-type Flag-tagged Hsmar1 transposase under the control of pEE (pRC1821, 1833 and 1845, negative control: pRC1806). **C/** Quantification of the number of papillae per colony. Average ± standard deviation of the mean of six representative colonies.

The EE-promoter-UTR sequence is not necessarily the highest level of activity attainable because transcription from the EE is likely stochastic and not every cell will have the same number of transcripts. Perhaps EE+ and EE++ are already in OPI when the cell has a single transcript due to a higher translation efficiency. We therefore explored transcriptional activity with a series of progressively degraded P_L_-λ promoters that had been selected from a mutant library for their lack of stochastic cell-to-cell variation (23).

### Characterization of the set of constitutive promoters

We synthesized a set of five constitutive promoters (00, JJ, K, E, and W) derived from the constitutive bacteriophage P_L_-λ promoter, based on (23). To increase the available range of expression levels, we also created a variant of each promoter where the RBS has been abolished. The expression construct is shown in Figure 3A and is composed of the promoter and a RBS sequence, NdeI and BamHI restriction sites facilitate cloning a gene of interest, which can then be fused or not to a C-terminal 3x FLAG tag. To avoid any read-through transcription, the construct is flanked by terminator sequences. The expression constructs were cloned either into a single-copy vector or a five-copy vector, pBACe3.6 and pGHM491, respectively. The following nomenclature will be used: Bp-EE to Bp6 represents the six promoters cloned into the single copy vector, Ip-EE to Ip6 corresponds to the six promoters cloned into the five copy vector, the ‘-‘ and ‘++’ represents the abolished or the optimized RBS, respectively.

**Figure 3.**
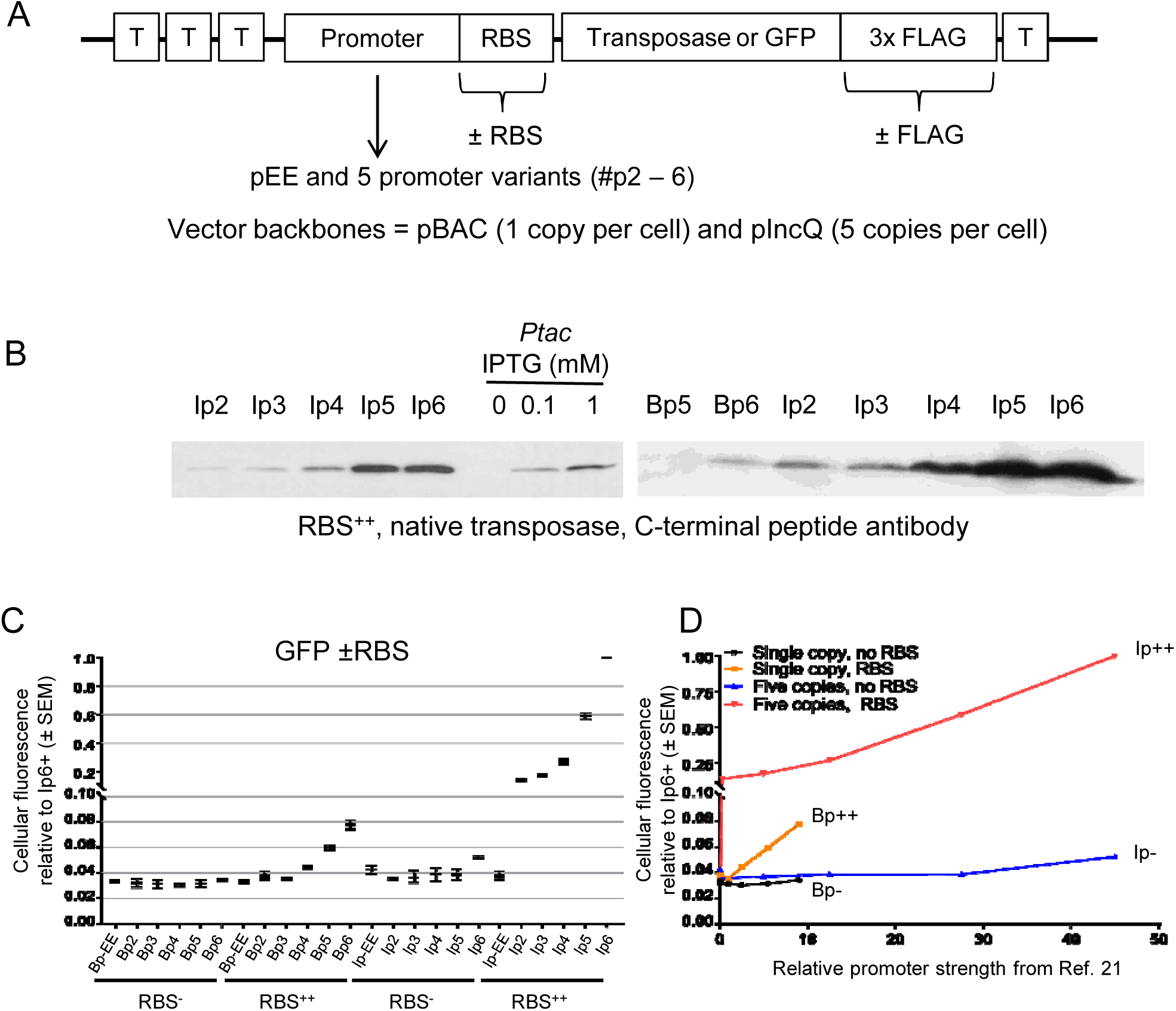
Characterization of the set of constitutive promoters. **A/** The *Hsmar1* gene is fused or not to 3x Flag tag on its C-terminus and cloned downstream of one of six different promoters (see text for more details) with an inactive or optimal RBS (defined in Figure 2A). The construct is located between terminator sequences (T) upstream and downstream to avoid read-through transcription. To further control the number of copies, the plasmid backbone is a one-copy, pBACe3.6, or a five-copy, pGMH491, vector. **B/** Western blots using an antibody against the C-terminus of Hsmar1, which compare the strongest promoters with an optimal RBS to the Ptac promoter induced with different concentration of IPTG. **C/** The promoter strength of each construct was determined by FACS after cloning an *EGFP* gene in each vector (pRC1782-1807). The number EE to 6 corresponds to one of the six promoters. The single and five-copy vectors are annotated B or I, respectively. The vectors with an inactive or an optimal RBS are annotated – or ++, respectively. Average ± standard deviation of the mean of three biological replicates. **D/** Plot of the average promoter strength (as defined in (23)) versus the promoter strength determined by FACS in Figure 3C.

We first investigated the strongest expression vectors by performing western blots with an anti-Hsmar1 antibody (Figure 3B). We also compared by western blotting these constructs with the Ptac inducible promoter previously used for papillation assay (Figure 3B). Interestingly, two of our constructs (Ip5++ and Ip6++) produce a higher amount of Hsmar1 transposase than the Ptac promoter fully induced with 1 mM of IPTG.

We next quantified the strength of each expression vector by inserting an *EGFP* gene in each Flag-tagged vector to investigate fluorescence levels by flow cytometry (Supplementary Figure 2). To rank the expression vectors, we normalized their average fluorescence value against the strongest vector, Ip6++ (Figure 3C). Most of the single-copy expression vectors produce an amount of EGFP fluorescence close to the detection threshold and therefore their ranking might not be accurate. However, all of the five-copy expression vectors produce more fluorescence than the single-copy vectors. Also, the vectors with a consensus RBS produce an amount of fluorescence that correlated with the promoter strength originally determined by Alper and colleagues (23). In contrast, all of the vectors without a RBS motif, except Ip6-, produce a fluorescence level close to the detection threshold (Figure 3D). Similarly, the pEE promoter is also too stochastic to change the amount of fluorescence produced whether the RBS is present or absent.

### Characterization of the papillation assay with the wild-type Hsmar1 transposase

To visually determine the best conditions for the papillation assay, we used the Ip3++ expression vector and a range of lactose concentrations (Supplementary Figure 3). We observed a correlation between the number of papillae per colony and the lactose concentration (Supplementary Figure 3A to C). We decided to work at 0.1% lactose since it represents the best trade-off between the number of papillae per colony and the size of the papillae for quantitation at high transposition rate.

We first investigated the transposition rate supported by each RBS^++^ expression vector with the wild-type transposase (Figure 4A). As expected from the wide range of expression, we observed a 350-fold variation in the average number of papillae per colony (Figure 4B). To better visualize the relationship between the expression vector strength and the transposition rate, as determined by the number of papillae per colony, we plotted the strength of the promoter as determined by Alper and colleagues (23) against the number of papillae per colony (Figure 4C). As previously documented *in vitro*, in *E. coli* and in HeLa cells, the wild-type Hsmar1 transposase follows an inverse-exponential relationship between transposase expression and transposition rate for Bp++ and Ip++ vectors (12, 21).

**Figure 4.**
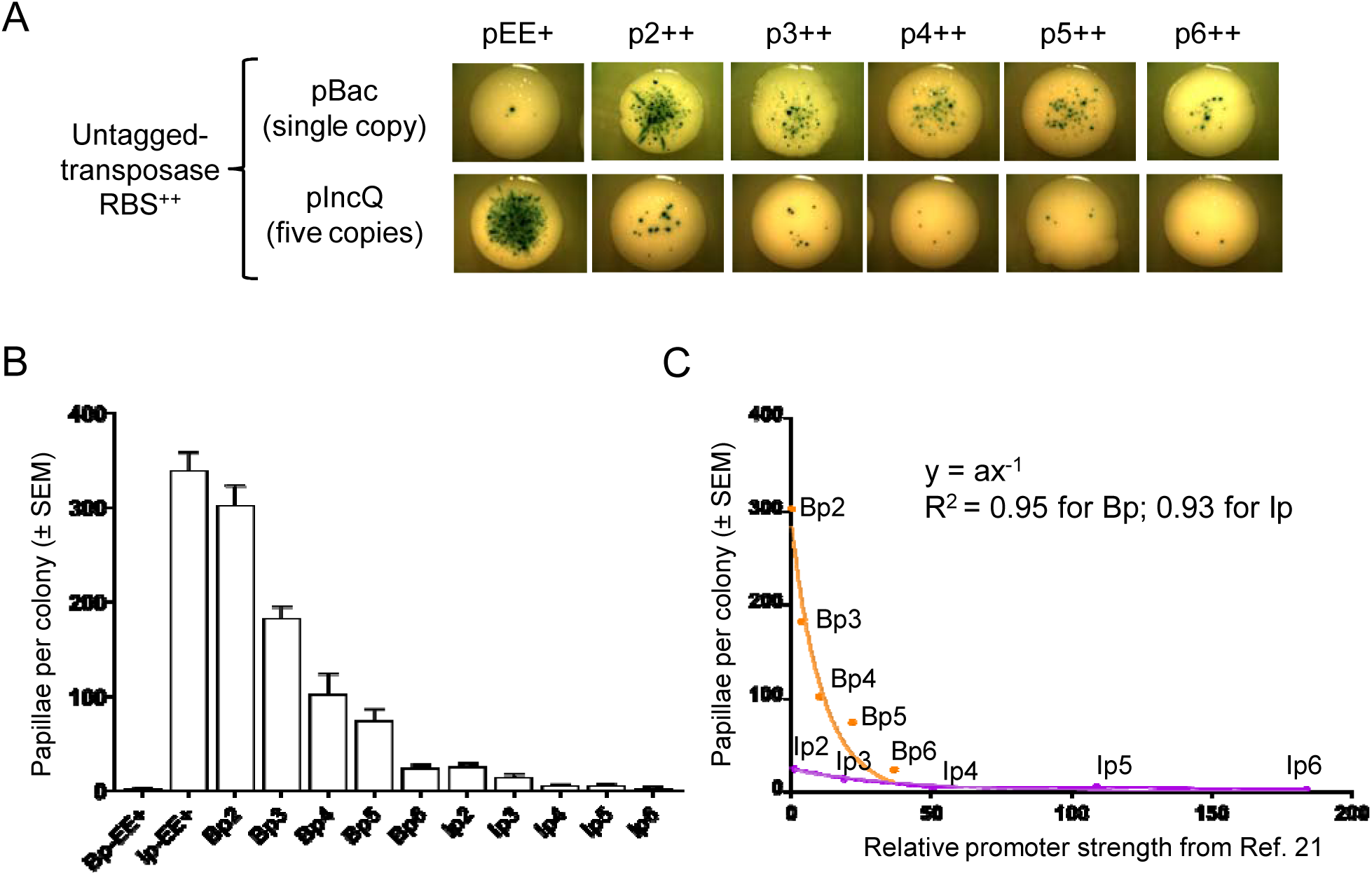
Characterization of the papillation assay with the wild-type untagged Hsmar1 transposase and optimal RBS. **A/** Representative colonies of each vector expressing a wild-type untagged Hsmar1 transposase (pRC1723-1728 and pRC1730-1735). **B/** Quantification of the number of papillae per colony. Average ± standard deviation of the mean of six representative colonies. **C/** Plot of the average promoter strength (as defined in (23)) versus the average number of papillae per colony (as defined in Figure 4B). As expected from overproduction inhibition (OPI), an inverse power law is observed between the promoter strength and the transposition rate.

There was a noticeable discontinuity in transposition rate between Bp5++ and Bp6++ and between pBac and pIncQ. We therefore tested the expression vectors with or without a RBS (Figure 5A). Quantitation of the transposition rate of each expression vector shows that the Bp++, Ip-, and Ip++ series follow an inverse-exponential relationship between transposase expression and transposition rate (Figure 5B). However, the set of Bp-expression vectors is more difficult to interpret because transcription and translation may be stochastic from cell to cell. This may be smoothed out in the Ip-series, which gave the most progressive response.

**Figure 5.**
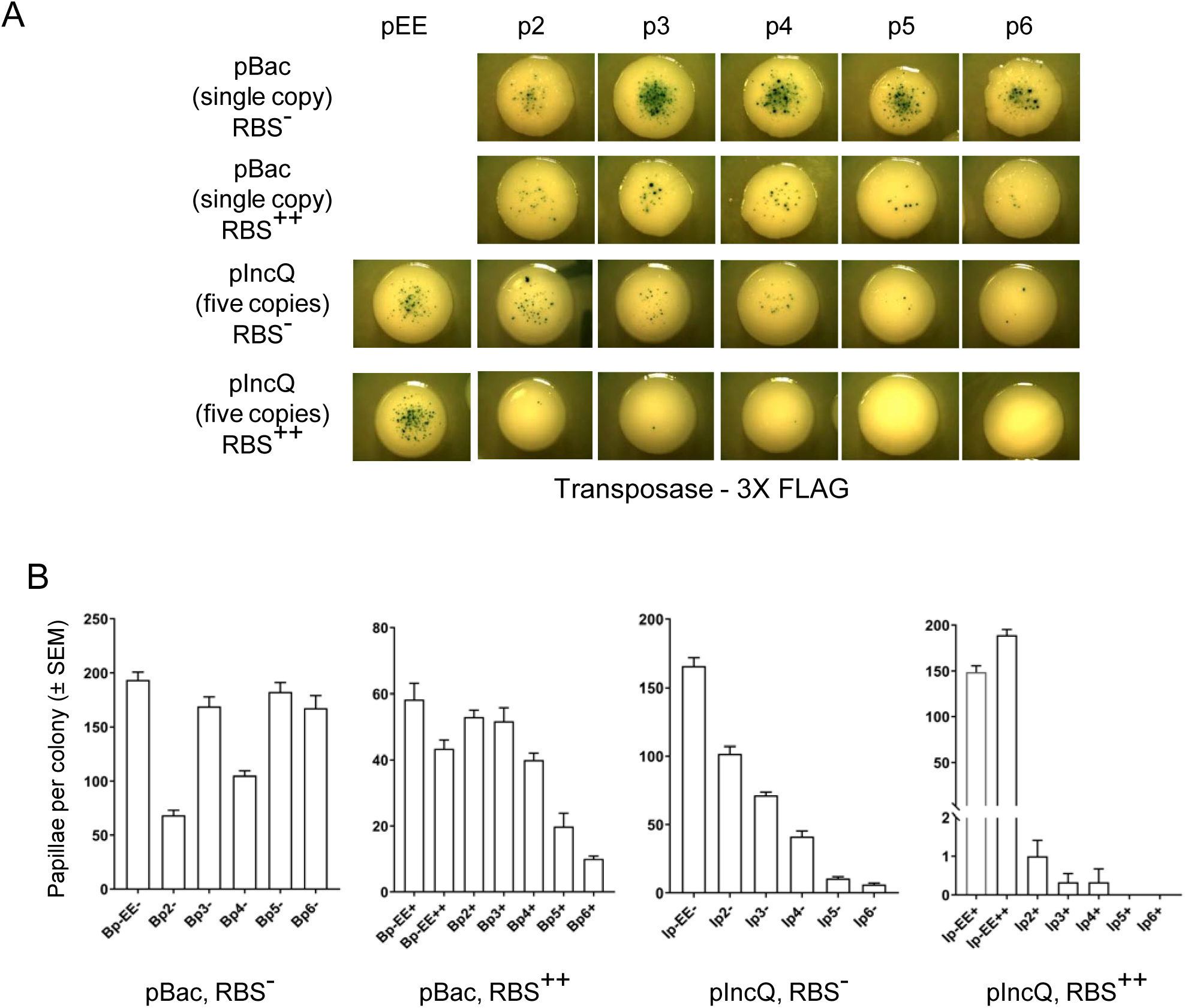
Characterization of the papillation assay with the wild-type Flag-tagged Hsmar1 transposase and an optimal or inactive RBS. **A/** Representative colonies of each vector expressing a wild-type Flag-tagged Hsmar1 transposase (pRC1821-1846). **B/** Quantification of the number of papillae per colony. Average ± standard deviation of the mean of six representative colonies.

For other transposons, the expression will have to be tuned to the system as different transposons will have different relationship between transposase concentration and transposition rate. A medium copy vector (pIncQ) with a medium promoter (p4) would be an ideal starting point. The expression can then be tuned by progressive degradation of the RBS.

### SETMAR transposition activity was lost during the same period as Hsmar1 transposase domestication

The Hsmar1 transposase was originally discovered in the human genome where an inactivated Hsmar1 transposase is fused to a SET domain to form the *SETMAR* gene (24-26). The domesticated Hsmar1 transposase is inefficient at performing transposition because of the mutation of the DDD triad catalytic motif to DDN (25, 26). We performed a papillation assay with an un-induced Ptac promoter driving expression of the D282N mutant derivative as well as 22 other mutations present in the human SETMAR to determine their effects on transposition (Figure 6A). Most of the mutations present in the human SETMAR are in the catalytic domain and are common to all anthropoid primates containing SETMAR, indicating that these mutations likely occurred before or during the domestication event. In addition to D282N, two other mutations, C219A and S279L, completely disrupt Hsmar1 transposition activity (Figure 6B). Two other mutations located in the DNA binding domain, E2K and R53C, also severely affect the transposition rate. In addition, seven other mutations located mostly in the catalytic domain mildly affect Hsmar1 transposition activity. Only one mutation, V201L, increases Hsmar1 transposition rate whereas the remaining mutations were neutral.

**Figure 6.**
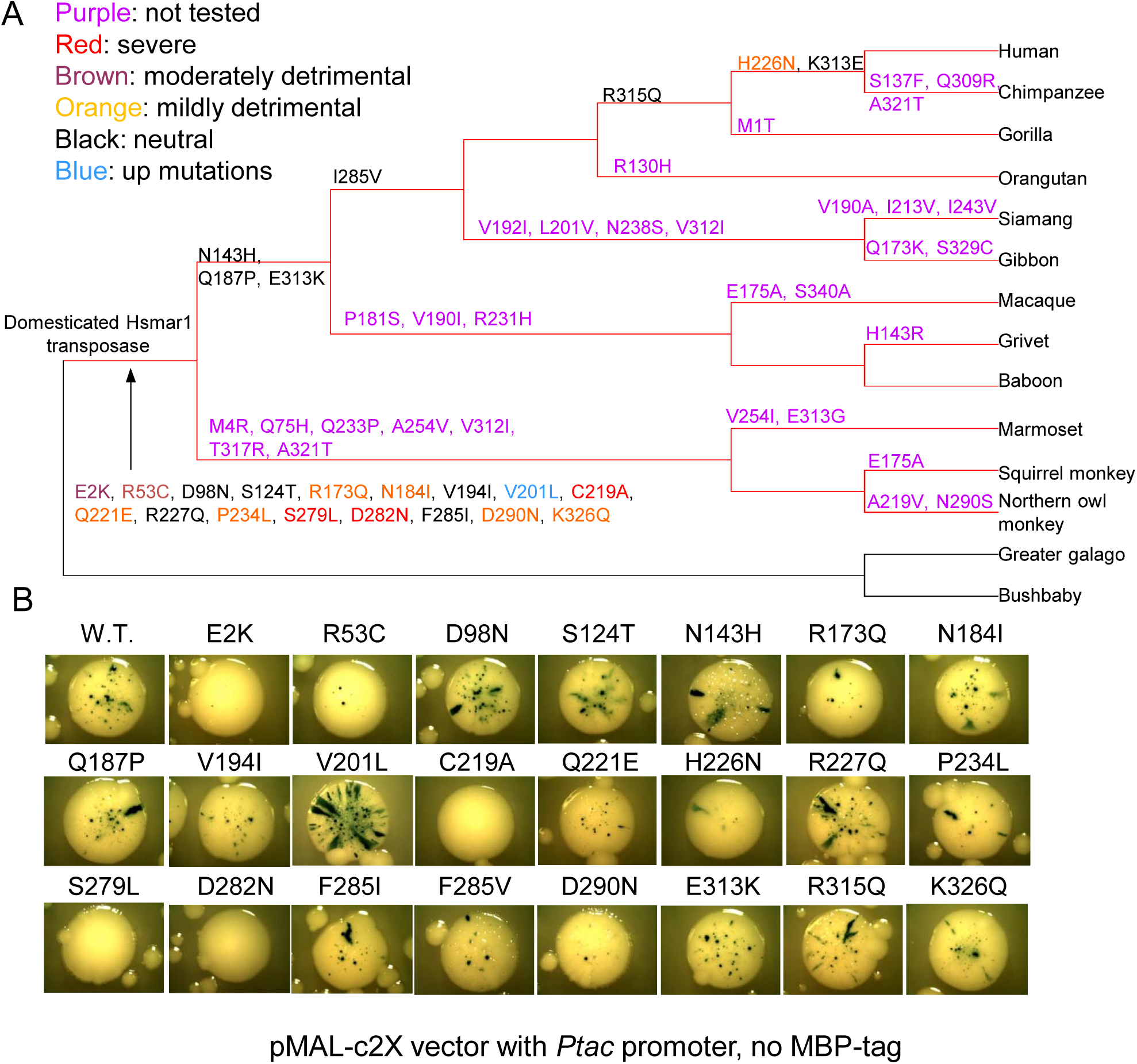
SETMAR transposition activity was lost during the same period as Hsmar1 transposase domestication. **A/** Phylogenetic tree of anthropoid primates which represents the apparition of mutations in the Hsmar1 domain of SETMAR. All the mutations present in the human SETMAR were tested by papillation assay to determine their effects on Hsmar1 transposition. **B/** Representative colonies of pMAL-C2X expressing wild-type (pRC1721) or mutant Hsmar1 transposases (pRC1877-1899).

Of the 23 mutations present in the Hsmar1 domain of SETMAR, 12 mutations are deleterious to the transposition rate, with three of them abolishing it completely (C219A, S279L and D282N). This result supports an absence of conservation of Hsmar1 nuclease activity during SETMAR evolution, in agreement with recent studies which did not observe an *in vivo* nuclease activity of SETMAR in DNA repair assays (27, 28). Two of the DNA binding mutants, E2K and R53C, are deleterious to Hsmar1 transposition activity in a papillation assay. It will be interesting to determine whether this effect is mediated through a change in ITR binding efficiency, which could have modified SETMAR’s ability to bind ITRs in the genome and therefore its emerging functions in regulating gene expression (29).

### Covalently linking two Hsmar1 monomers in a dimer affects the transposition rate

We recently described a novel Hsmar1 transposase construct where two monomers are covalently bound by a linker region (30). We took advantage of our approach to test whether the transposition rate of a covalently bound Hsmar1 dimer differs from that of the Hsmar1 monomer. At low expression levels, we expect a covalently bound Hsmar1 dimer to transpose more efficiently than an Hsmar1 monomer because of the physical link between the subunits, which favors dimerization and also requires only a single translation event. We cloned the monomeric and dimeric construct in a set of expression vectors spanning very low to high expression and performed a papillation assay (Figure 7A). In agreement with our model, we observe a change in the number of papillae per colony with vectors with the lowest expression levels, as shown by the quantitation in Figure 7B.

**Figure 7.**
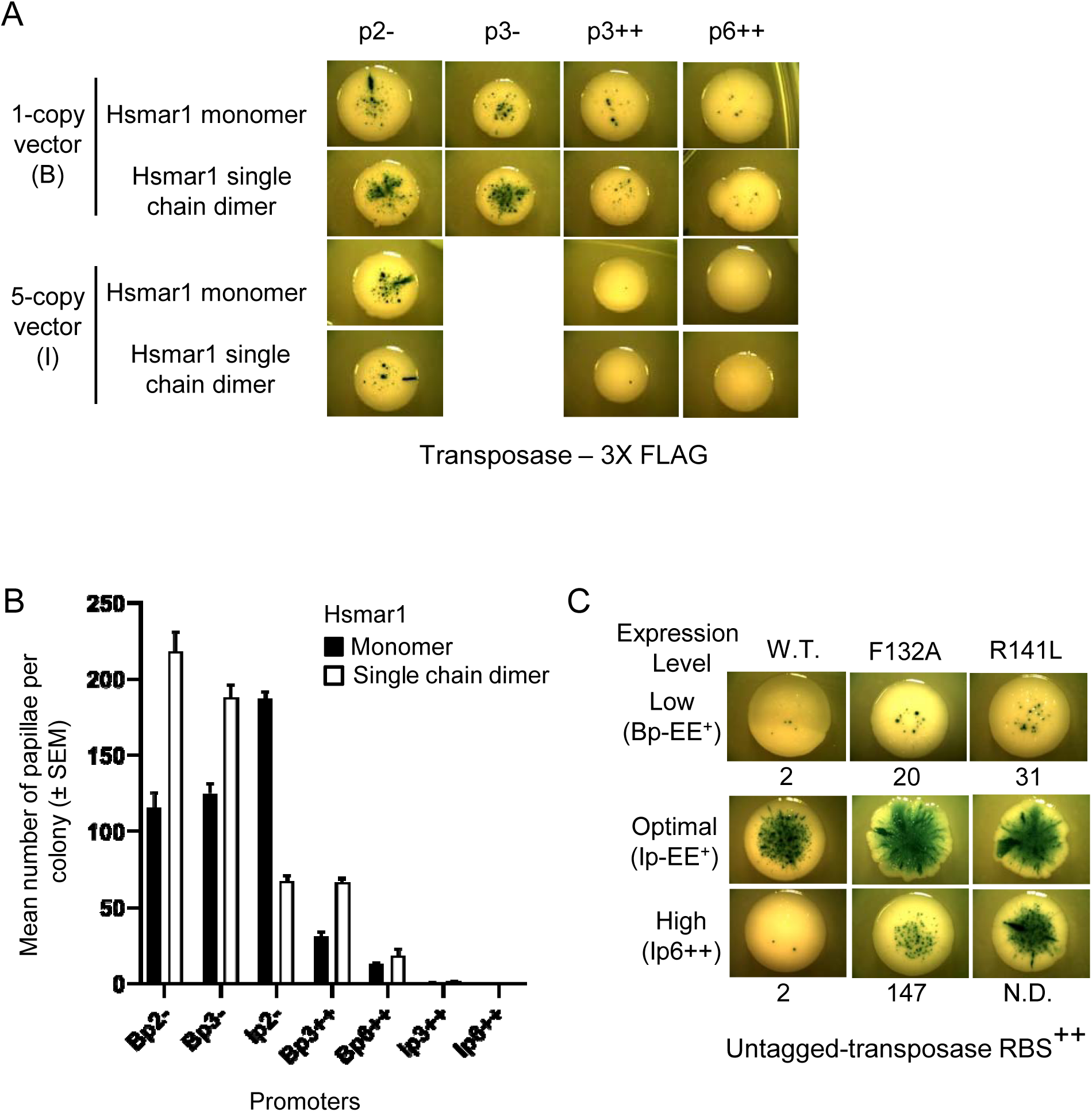
Covalently linking two Hsmar1 monomers in a dimer or mutating Hsmar1 dimer interface affect the transposition rate. **A/** Representative colonies of each expression vector expressing either Hsmar1 monomer (pRC1868-1871, 1873, 1875, and 1876) or Hsmar1 single chain dimer (pRC1858-1861, 1863, 1865, and 1866). **B/** Quantification of the number of papillae per colony. The expression vectors have been ordered by decreasing number of papillae per colony for the Hsmar1 monomer. Average ± standard deviation of the mean of six representative colonies. **C/** Different Hsmar1 mutants have been tested in low, optimal and high transposase expression level (Bp1+ (pRC1739 and 1740), Ip1+ (pRC1746 and 1747) and Ip6++ (pRC1752 and 1753), respectively). Representative colonies of each papillation plate is shown. The average number of papillae per colony is indicated below the pictures. Average ± standard deviation of the mean of six representative colonies.

When compared to the results obtained with Hsmar1 monomer, the covalent dimer transposition rate peaks at a different set of expression vectors, Bp2- and Bp3-for the covalent dimer and Ip2-for the monomer (Figure 7B). These three expression vectors have a similar relative promoter strength, around 4% of Ip6++ (Figure 3C), indicating that the number of transposases molecules expressed per cell is particularly low. Based on this idea, we can hypothesize that Bp2- and Bp3-, which provide the highest transposition rates for the single chain dimer, are weaker promoters than Ip2-, which provides the highest transposition rate for the monomeric Hsmar1 but a lower transposition rate for the single chain dimer. Thus, Bp2- and Bp3- are likely to express on average less than two proteins per cell, which is not sufficient to promote optimal transposition for the Hsmar1 monomer construct. In contrast, Ip2- is likely to express on average at least two proteins per cell, which starts to promote OPI for the covalent dimer construct and therefore results in a lower transposition rate than Bp2- and Bp3-. Inversely, we do not observe any difference in the number of papillae per colony with stronger expression vectors such as Ip3++ and Ip6++ (Figure 7A and B). This indicates that a covalently bound Hsmar1 dimer is as sensitive to OPI as the Hsmar1 monomer.

### Mutations in Hsmar1 dimer interface produce hyperactive mutants in bacteria

The mutagenic nature of transposable elements makes them useful in screening for essential genes. However, OPI limits the transposition rate when the transposase concentration is too high (12). One way to overcome OPI is to decrease the stability of the Hsmar1 dimer to shift the monomer-dimer equilibrium to the inactive monomeric form. We decided to take advantage of our approach to investigate two Hsmar1 transposases mutated in the dimer interface, one known mutant, F132A (F460 in SETMAR (31)), and a novel one, R141L (9). We used three vectors expressing Hsmar1 transposase at a low (Bp-EE+), optimal (Ip-EE+), and high (Ip6++) expression level (Figure 7C). The average number of papillae per colony is indicated below each representative colony. Interestingly, both F132A and R141L transposases are hyperactive at low and optimal levels of expression when compared to WT. A higher transposition rate is also observed at high expression level for both mutants, with R141L showing a stronger resistance to OPI than F132A. To confirm the results, the transposition rates were also determined using the mating-out assay (19), which is more quantitative (Table 1). The results of the mating-out and transposition assays were similar. Interestingly, Hsmar1 R141L transposition rate is not affected by the high transposase expression level produced by Ip6++, as the rate remains similar between Ip- EE+ and Ip6++ whereas we observe a 147-fold and a 17-fold decrease for the wild type transposase and for the F132A mutant, respectively.

**Table 1:**
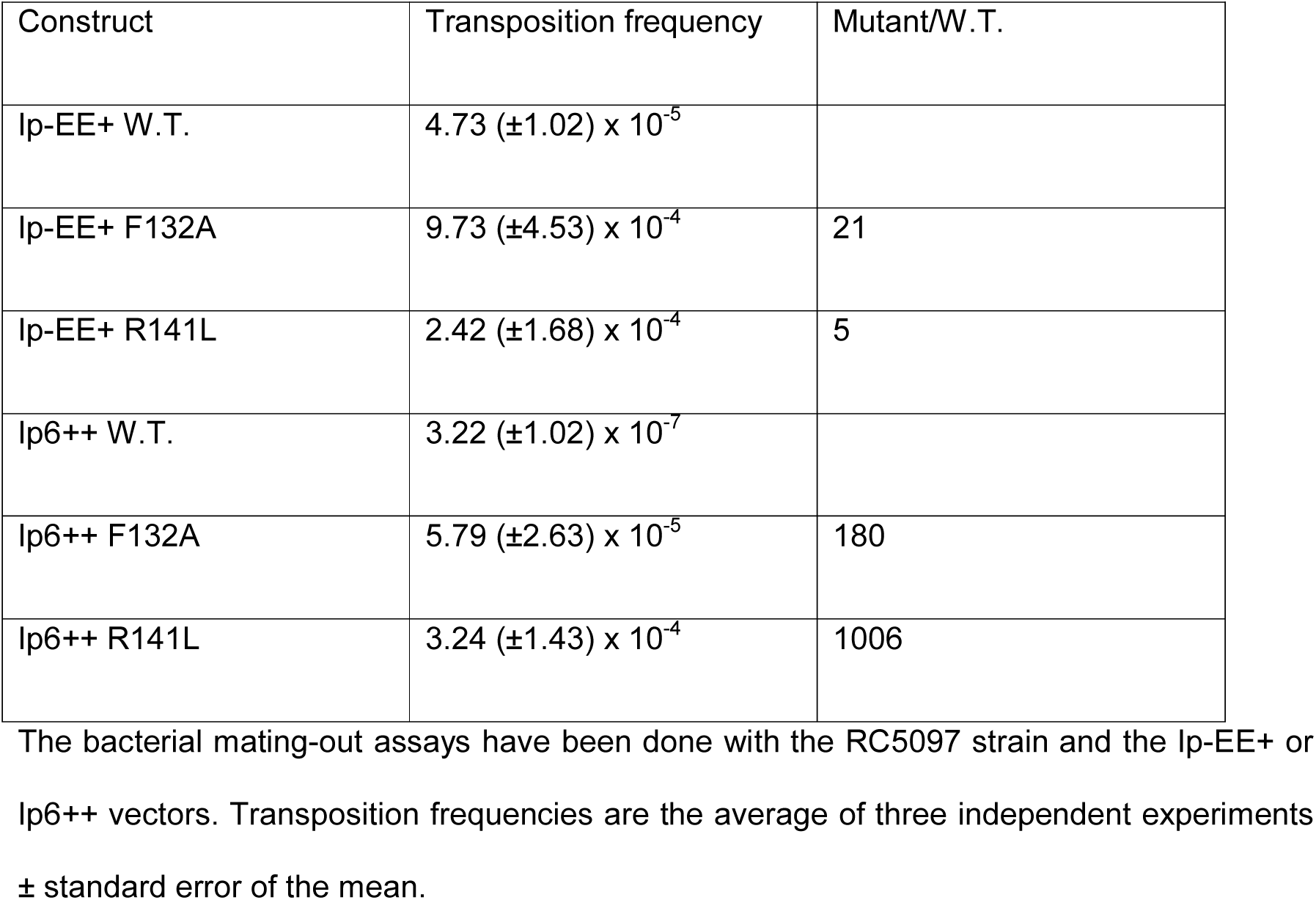
Transposition frequencies of two Hsmar1 transposase mutants expressed at optimal and high level.

The hyperactivity of F132A and R141L mutants could be explained by the promotion of one or more of the conformational changes during the reaction (11). The decreased OPI-sensitivity could result from a decrease in the dimer stability, which shifts the monomer-dimer equilibrium towards the monomeric form, and therefore reduces the concentration of active transposases in the cell. Also, an unstable dimer bound to a transposon end could be more likely to fall apart allowing the recruitment of the previously bound end by another bound dimer, activating transposition. This type of mutant is more likely to exhibit hyperactivity only in bacteria. Indeed, in mammalian cells the size of the nucleus and the larger quantity of non-specific DNA would be expected to increase the time necessary for a transposase to find a transposon end (21). Therefore, transposases with a weakened dimer interface are more likely to revert to an inactive monomeric state resulting in hypoactive mutants.

## Conclusion

This study provides a set of expression vectors based on constitutive promoters to investigate the phenotypes of mutant transposase. It will be useful to distinguish between true hyperactive mutants and defective mutants that happen to be resistant to OPI. Compared to inducible promoters, our set of expression vectors provides a wide range of consistent transposase expression levels between individual cells. In addition to the characterization of the constitutive promoters, we also found one Hsmar1 mutation, R141L, which is OPI-resistant in *E. coli* and could therefore prove useful for improving bacterial transposon mutagenesis with *mariner* elements. Another approach in controlling the transposition rate is to covalently bind two Hsmar1 monomers, which allows transposition to occur after a single translation event and therefore permits the usage of a weak promoter with a weak RBS.

We believe our set of expression vectors will be useful or the study of other transposons and in the screening of libraries for finding hyperactive and/or OPI-resistant transposases.

## Methods

### Media and bacterial strains

Bacteria were grown in Luria-Bertani (LB) media at 37°C. The following antibiotics were used at the indicated concentrations: ampicillin (Amp), 100 µg/ml), chloramphenicol (Cm), 25 µg/ml, and spectinomycin (Spec), 100 µg/ml. The following *E. coli* strains were used: RC5024 (identical to DH5α) [endA1 hsdR17 glnV44 thi-1 recA1 gyrA relA1 Δ(lacIZYA-argF)U169 deoR (φ80dlac Δ(lacZ)M15)], RC5094 [F-araD139 Δ(argF-lac)U169 rspL150 relA1 flbB5301 fruA25 deoC1 ptsF25 rpoS359::Tn*10*], RC5096 [F^-^ fhuA2 Δ(lacZ)r1 glnV44 e14-(McrA-) trp-31 his-1 rpsL104 xyl-7 mtl-2 metB1 Δ(mcrC-mrr)114::*IS10* argE::Hsmar1-lacZ’-kanR] and RC5097 (= RC5096 pOX38::miniTn10-CAT).

### Constitutive promoters

Alper et al previously generated and characterized a set of constitutive promoters based on P_L_-λ ranging from strong down to very weak (23). We select the promoters 00, jj, K, E, and W (equivalent to p2, p3, p4, p5, and p6 in this study) and generate pEE, a featureless tract of 44 GACT repeats which we represent an ideal promoter-less region (Table 2). Each promoter sequence is preceded by three terminator sequences and followed by a consensus ribosome binding site (RBS++), a null RBS (RBS-), or a GACT RBS in the case of pEE (RBS+), a transposase gene, three Flag tag and a terminator sequence (Figure 2A).

**Table 2:**
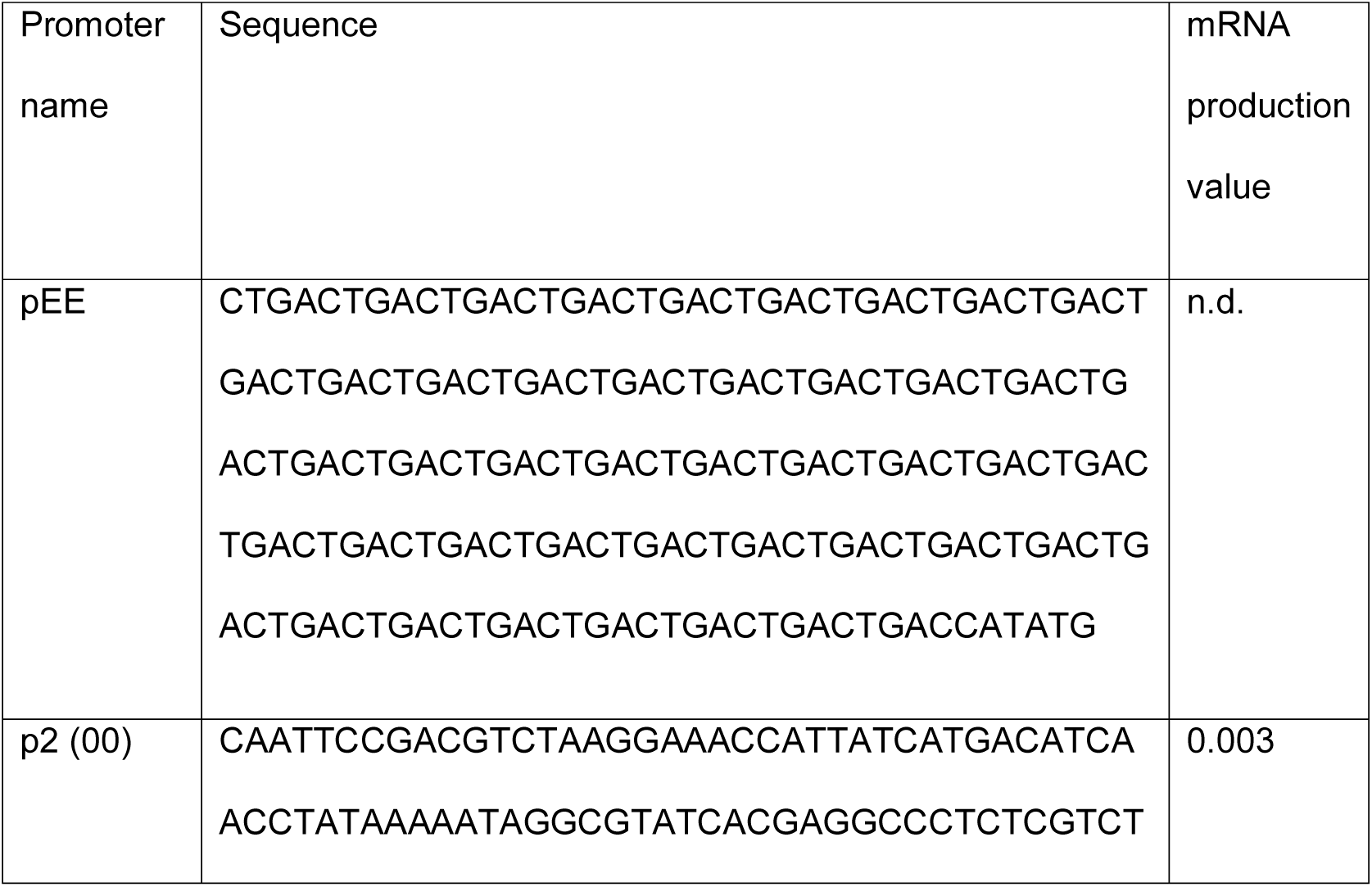
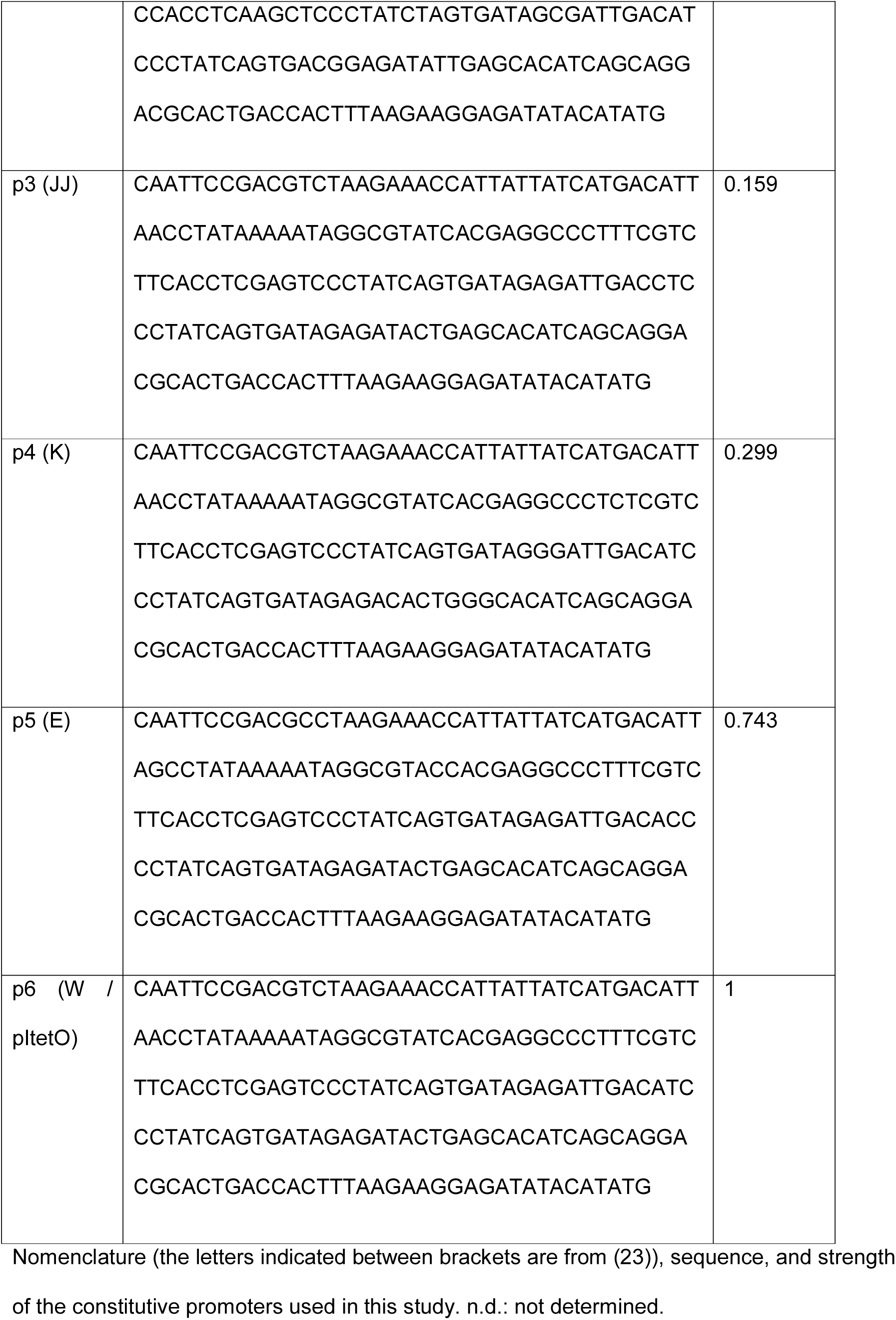
List of constitutive promoters.

### Plasmids

Expression plasmids were built by cloning the *EGFP* or *Hsmar1* gene in pBACe3.6, pGHM491, and pMAL-c2X (New England Biolabs) between NdeI and BamHI restriction endonuclease sites. A list of the plasmids used in this study can be found in Supplementary Table 1. The DNA sequences of the vectors based on pBACe3.6 and pMAL-c2X can be found in Supplementary Table 2. The DNA sequence of pGHM491 is unknown and therefore the DNA sequences of the vectors based on it are absent from Supplementary Table 2. Plasmids pRC880 and pRC1721 encode the wild-type transposase in pMAL-c2X in presence and absence of the MBP tag, respectively (Figure 1). Plasmids pRC1782-1807 encode EGFP downstream of pEE to p6, with RBS-, RBS+, and RBS++, in pBACe3.6 and pGHM491 (Figure 3). Plasmids pRC1723-1728 and pRC1730-1735 encode untagged Hsmar1 downstream of pEE to p6, with RBS+ and RBS++, in pBACe3.6 and pGHM491 (Figures 2 and 4). Plasmids pRC1821-1846 encode Flag-tagged Hsmar1 downstream of pEE to p6, with RBS-, RBS+, and RBS++, in pBACe3.6 and pGHM491 (Figures 2 and 5). Plasmids pRC1877 to pRC1899 are derived from pMAL-c2X and encode the different Hsmar1 mutants with the mutations found in SETMAR (Figure 6). Plasmids pRC1858-1861, 1863, 1865, 1866, 1868-1871, 1873, 1875, and 1876 encode the Hsmar1 monomer and Hsmar1 single chain dimer in Bp2-, Bp3-, Bp3++, Bp6++, Ip2-, Ip3++, and Ip6++ (Figure 7). Plasmids pRC1739, 1740, 1746, 1747, 1752, and 1753 encode Hsmar1 F132A and R141L mutants cloned into Bp-EE+, Ip-EE+, and Ip6++ (Figure 7).

### Flow cytometry

RC5096 cells expressing EGFP were grown overnight at 37°C in LB medium supplemented with chloramphenicol or spectinomycin. The cultures were diluted in a 1:1000 ratio in fresh LB medium complemented with antibiotics and grown to mid-log phase (OD_600_ ∼ 0.5). The cells were pelleted at 6,000g for 5 min, washed in 1X PBS twice, and resuspended in 500 µl of 1X PBS. Flow cytometry analysis was performed on 100,000 cells with a Beckman Coulter Astrios EQ and data analysed using Weasel software v3.0.2.

### Western blotting

Cells containing a derivative of pMAL-c2x were grown in LB supplemented with 100 μg/ml of ampicillin at 37°C until an OD_600_ of ∼ 0.5 and were then induced with the required concentration of IPTG for 2 hours at 37°C. Cells containing pGHM491 or pBACe3.6 derivatives were grown in LB supplemented with respectively 100 μg/ml of spectinomycin or 50 μg/ml of chloramphenicol at 37°C for the same amount of time as the induced cells. Promoters’ expression was analysed by pelleting ∼1.5×10^9^ cells. The samples were resuspended in SDS sample buffer, boiled for 5 min, and loaded on 10% SDS-PAGE gels. Proteins were transferred to PVDF membrane, probed with an anti-Hsmar1 antibody (goat polyclonal, 1:500 dilution, ab3823, Abcam) followed by a horseradish peroxidase-conjugated anti-goat secondary antibody (rabbit polyclonal, 1:5000 dilution, ab6741, Abcam). Proteins were visualized by using the ECL system (Promega) and Fuji medical X-ray film (Fujufilm).

### Papillation assay

The papillation assay and the reporter strain RC5096 have been described previously (Supplementary Figure 1) (18). Briefly, transposase expression vectors were transformed into the RC5096 strain. It is a lac^-^ *E. coli* strain encoding a transposon containing a promoter-less lacZ and a kanamycin resistance gene flanked with Hsmar1 ends, which has been integrated in a silent genomic locus. In absence of LacZ, the strain produces white colonies on X-gal indicator plates. When the transposase is supplied *in trans*, the integration of a transposon into the correct reading frame of an active gene will produce a lacZ fusion protein. The descendants of this cell will become visible as blue papillae on X-gal indicator plates. RC5096 transformants were plated on LB-agar medium supplemented with 0.01% lactose, 40 μg/ml of X-gal and either 50 μg/ml of chloramphenicol or 100 μg/ml of spectinomycin. Plates were incubated 5 days at 37°C and photographed. The transposition rate is determined by the number of papillae per colony. Papillation assays were performed in biological duplicates.

### Mating-out assay

A chloramphenicol resistant derivative of the conjugative plasmid pOX38 has been introduced in the RC5096 papillation strains to create the donor strains RC5097. Briefly, RC5097 transformants and the recipient strain, RC5094, were grown overnight in LB supplemented with antibiotics at 37°C. The next day, respectively one and three volumes of RC5097 and RC5094 were centrifuged for 5 min at 6,000x g. Each pellet was resuspended in 3 ml of fresh LB, pool together, and incubated in a shaking water bath for 3 hours at 37°C. After the mating, the transposition events were detected by plating 200 μl of each culture on LB-agar medium supplemented with tetracycline and kanamycin. The number of transconjugants was obtained by plating a 10^-5^ fold dilution of each culture on LB-agar medium supplemented with tetracycline and chloramphenicol. The plates were incubated overnight at 37°C and the transposition rate determined the next day by dividing the number of kanamycin-resistant cells by the number of chloramphenicol resistant cells.

## List of abbreviations

EE: “even-end” promoter
ITR: inverted terminal repeat
OPI: overproduction inhibition
RBS: ribosome binding site
TE: transposable element.

## Acknowledgments

We would like to thank Michael Chandler for his comments on the manuscript. We also thank David Onion from the University of Nottingham Flow Cytometry facility for help with FACS analyses.

## Funding

This work was supported by the Wellcome Trust [WT093160] to RC and a Biotechnology and Biological Sciences Research Council Doctoral Training Program Grant [BB/J014508/1] to MT.

## Availability of data and materials

All the materials mentioned and used in this work will be made available upon request.

## Authors’ contributions

Performed the experiments: MT. Conceived and designed the experiments and analysed the data, MT, RC. Wrote the paper: MT, RC. All Authors read and approved the final version the manuscript.

## Ethics approval and consent to participate

Not applicable.

## Consent for publication

Not applicable.

## Competing interests

The Authors declare that they have no competing interests.

