## Supplementary Figures for "A series of constitutive expression vectors to accurately measure the rate of DNA transposition and correct for auto-inhibition"

### Slide 1
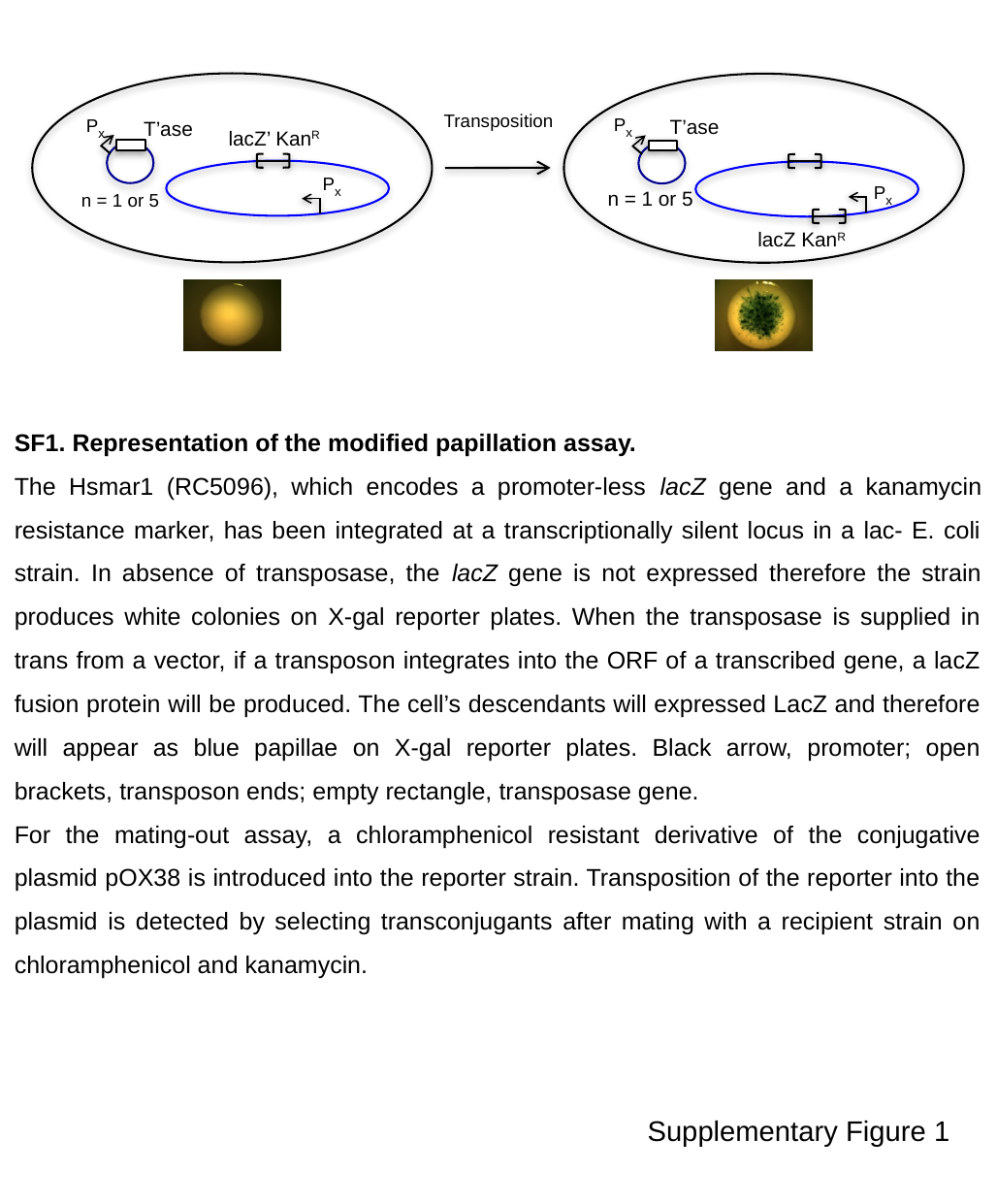

P_x_
Transposition
P_x_
T’ase
T’ase
lacZ’ Kan^R^
P_x_
P_x_
n = 1 or 5
n = 1 or 5
lacZ Kan^R^
SF1. Representation of the modified papillation assay.
The Hsmar1 (RC5096), which encodes a promoter-less lacZ gene and a kanamycin resistance marker, has been integrated at a transcriptionally silent locus in a lac- E. coli strain. In absence of transposase, the lacZ gene is not expressed therefore the strain produces white colonies on X-gal reporter plates. When the transposase is supplied in trans from a vector, if a transposon integrates into the ORF of a transcribed gene, a lacZ fusion protein will be produced. The cell’s descendants will expressed LacZ and therefore will appear as blue papillae on X-gal reporter plates. Black arrow, promoter; open brackets, transposon ends; empty rectangle, transposase gene.
For the mating-out assay, a chloramphenicol resistant derivative of the conjugative plasmid pOX38 is introduced into the reporter strain. Transposition of the reporter into the plasmid is detected by selecting transconjugants after mating with a recipient strain on chloramphenicol and kanamycin.
Supplementary Figure 1

### Slide 2
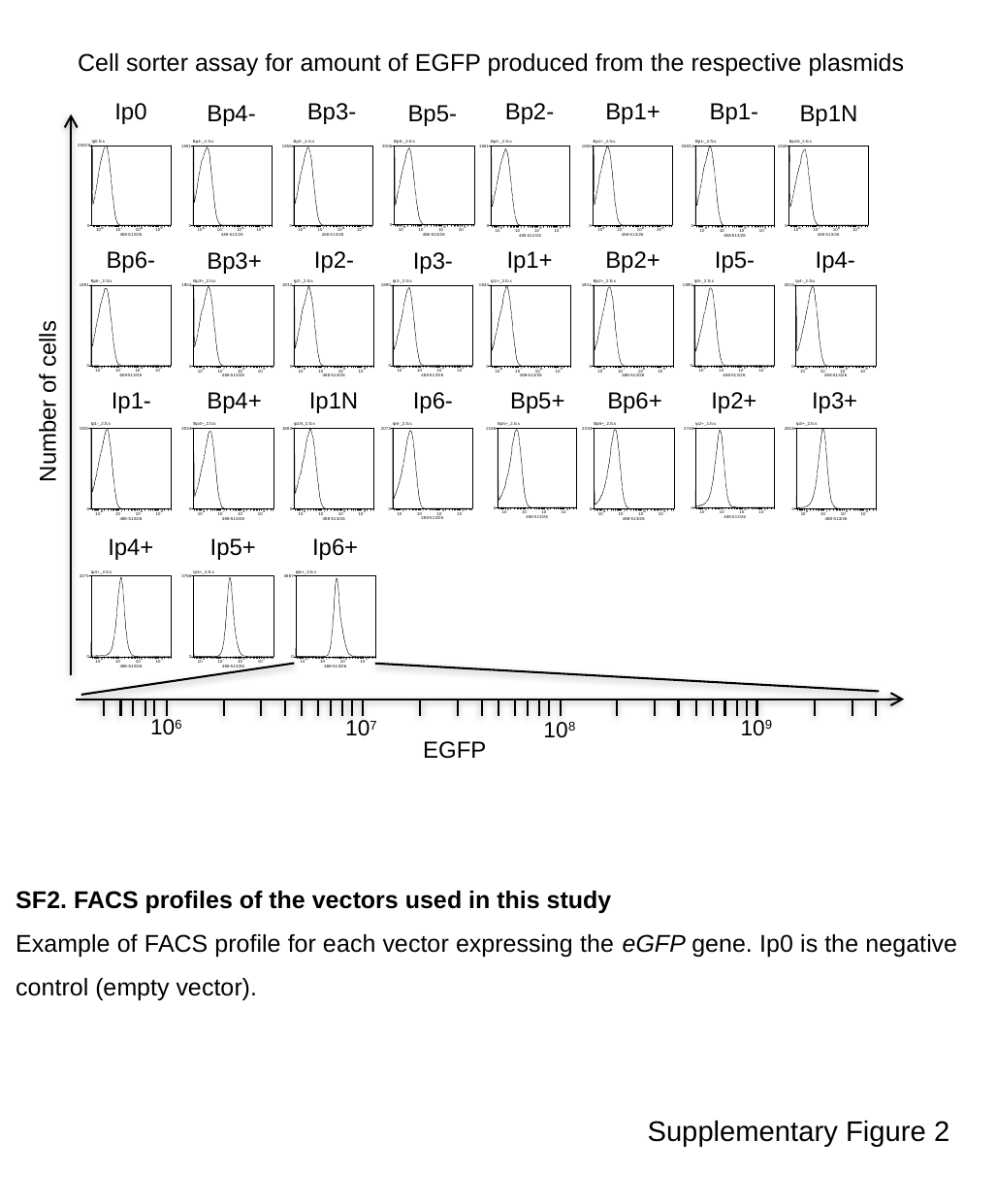

Cell sorter assay for amount of EGFP produced from the respective plasmids
Ip0
Bp3-
Bp2-
Bp1+
Bp1-
Bp4-
Bp5-
Bp1N
Bp6-
Ip2-
Ip1+
Bp2+
Ip5-
Ip4-
Bp3+
Ip3-
Ip1-
Bp4+
Ip1N
Ip6-
Bp5+
Bp6+
Ip2+
Ip3+
Number of cells
Ip4+
Ip5+
Ip6+
10^6^
10^9^
10^7^
10^8^
EGFP
SF2. FACS profiles of the vectors used in this study
Example of FACS profile for each vector expressing the eGFP gene. Ip0 is the negative control (empty vector).
Supplementary Figure 2

### Slide 3
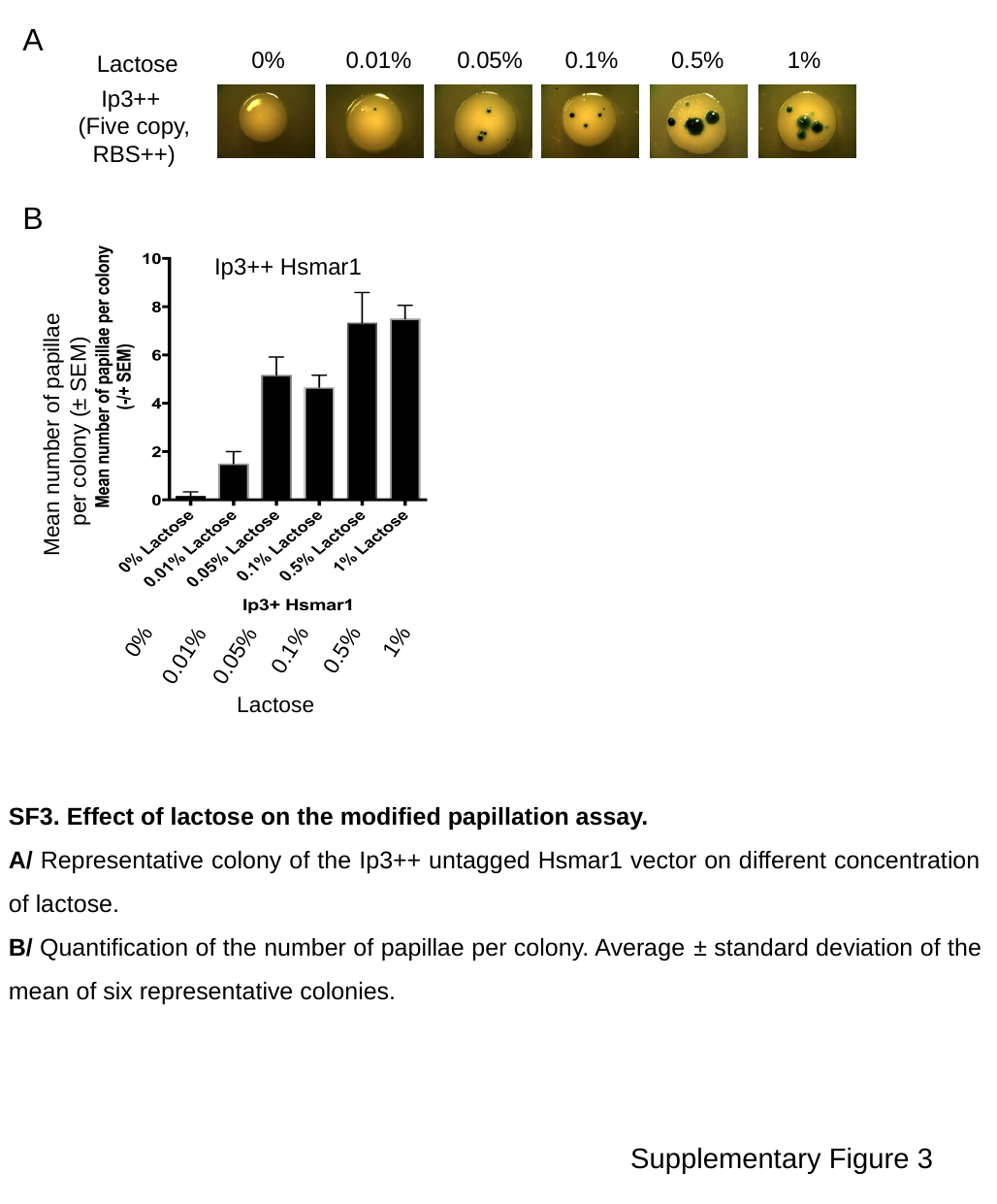

A
0%
0.01%
0.05%
0.1%
0.5%
1%
Lactose
Ip3++
(Five copy,
RBS++)
B
Ip3++ Hsmar1
Mean number of papillae
per colony (± SEM)
0%
1%
0.1%
0.5%
0.01%
0.05%
Lactose
SF3. Effect of lactose on the modified papillation assay.
A/ Representative colony of the Ip3++ untagged Hsmar1 vector on different concentration of lactose.
B/ Quantification of the number of papillae per colony. Average ± standard deviation of the mean of six representative colonies.
Supplementary Figure 3
